# A multistage, dual voxel study of glutamate in the anterior cingulate cortex in schizophrenia supports a primary pyramidal dysfunction model of disorganization

**DOI:** 10.1101/2023.11.27.568930

**Authors:** Lejia Fan, Zhenmei Zhang, Xiaoqian Ma, Liangbing Liang, Yujue Wang, liu Yuan, Lijun Ouyang, Zongchang Li, Xiaogang Chen, Ying He, Lena Palaniyappan

## Abstract

**Background:** Schizophrenia is an illness where glutamatergic dysfunction in the anterior cingulate cortex (ACC) has been long suspected; Recent in vivo evidence (Adams et al. 2022) has implicated pyramidal dysfunction (reduced glutamate tone) as the primary pathophysiology contributing to subtle features, with a secondary disinhibition effect (higher glutamate tone) resulting in the later emergence of prominent clinical symptoms. We investigate if genetic high risk (GHR) for schizophrenia reduces glutamatergic tone in ACC when compared to the states of clinical high risk (CHR) and first episode schizophrenia (FES) where symptoms are already prominent.

**Methods:** We recruited 302 individuals across multiple stages of psychosis (CHR, n=63; GHR, n=76; FES, n=96) and healthy controls (n=67) and obtained proton magnetic resonance spectroscopy of glutamate from perigenual ACC (pACC) and dorsal ACC (dACC) using 3-Tesla scanner.

**Results:** GHR had lower Glu compared to CHR while CHR had higher Glu compared to FES and HC. Higher disorganization burden, but not any other symptom domain, was predicted by lower levels of Glu in the GHR group (dACC and pACC) and in the CHR group (pACC only).

**Conclusions:** The reduction in glutamatergic tone in GHR supports the case for a pyramidal dysfunction contributing to higher disorganization, indicating disorganization to be the core domain in the pathophysiology of schizophrenia. Higher glutamate (likely due to disinhibition) is apparent when psychotic symptoms are raising to be prominent (CHR), though at the full-blown stage of psychosis, the relationship between glutamate and symptoms ceases to be a simple linear one.

## 1. Introduction

Schizophrenia is an illness where glutamatergic dysfunction in anterior cingulate cortex (ACC) has been long suspected; but in vivo studies of glutamate in early stages of this illness (high-risk states) have been inconclusive. There is no consensus on whether glutamate levels are consistently increased or decreased in patients(1–5); it is also unclear whether glutamatergic changes are of direct relevance to core symptoms of schizophrenia (6–9) or if they only index a genetic susceptibility (10–13). These questions are important for validating in vivo the prevalent microcircuit level models of schizophrenia centered on pyramidal dysfunction and excitotoxicity.

One of the most compelling recent accounts of glutamate dysfunction and phenotype of schizophrenia is the account put forth by Adams et al (14). Using an electrophysiological data based modelling of circuit level dysfunction, Adams et al. argued that a primary pyramidal dysfunction (reduced glutamate tone) is at the forefront of the pathophysiology of schizophrenia. However, expression of clinical symptoms such as hallucinations and delusions rely on a secondary disinhibition effect (higher glutamate tone) that results from a downstream (mal)adaptive change in inhibitory feedback. This argument suggests that individuals with genetic predisposition for schizophrenia will have a low glutamatergic tone (indexed by reduced glutamate in the ACC (11–13), when compared to those who express either subthreshold or full-blown symptoms of schizophrenia where symptoms are already apparent. Extending this further, we can expect low glutamate levels (as the primary phenomenon) to relate to a core, phenotype of the illness reflecting the primary effect of pyramidal dysfunction. We argue for this core feature that is subtle but still detectable in early pre-psychotic stages to be the primary symptom domain of disorganization. As illness develops into an incipient psychotic break with a subthreshold symptom complex (i.e., clinical high risk state), the secondary disinhibition effect will be reflected in a relative glutamate excess; at this stage, the primary symptom domain is still apparent but the secondary (positive symptoms) are emerging (14–16). When the state of disinhibition is well established (i.e. normalization of initial deficit), psychosis will be fully expressed as high levels positive symptoms (first episode psychosis) and glutamate levels will be normalized and be comparable to healthy people.

The model of relating a subtle neurocognitive symptom domain to genetic liability follows the framework laid out by Paul Meehl, who described ‘cognitive slippage’ (or loosened associations) to be a core feature of the integrative defect (disorganization) that emerges from an “aberration of the synaptic control system”(17). Several lines of evidence since Meehl’s proposition place disorganization (i.e., cognitive features of poor attention, poor abstract thinking, inappropriate or odd speech, behavior and affect) as the candidate symptom domain tracking the primary pathophysiology of schizophrenia. Unlike the symptoms of reality distortion that are episodic and emerge later in the course, disorganized thinking is seen many years before the first psychotic episode in those at risk (by mid-childhood/adolescence (18), often in conjunction with impoverished mental activity (together called ‘Bleulerian factor’ by Dominguez et al. (19) and the core of classical schizophrenia by Liddle et al. (20)). The genetic diathesis to schizophrenia, reflected in the continuous polygenic risk scores of this illness, map on to the burden of disorganization symptoms more than any other symptom domain (including negative symptoms) (21). Several studies have found that the cognitive deficits that are pervasive and influential in determining overall outcomes of schizophrenia track the severity of disorganization more closely than other symptom domains (22–26). Persistent disorganization is one of the most influential factors determining poor employment and educational outcomes in psychosis (27, 28). Demjaha and colleagues first demonstrated that among the various symptom dimensions, the burden of disorganization is the best predictor of the onset of full blown psychosis in those at CHR state (29); several clinical ratings (30) and speech-based studies later confirmed this with objective ratings (31, 32), supporting our argument that primary pathophysiological process that predates full-blown psychosis must relate to disorganization. Furthermore, after several acute episodes, the symptom profile in those with established schizophrenia shows “a drift toward disorganization (hebephrenia) (33)”, adding further support for disorganization being a candidate domain that tracks putative primary pathophysiology of schizophrenia.

To test the proposition that the primary pathophysiology is characterized by low glutamatergic tone evident in the genetically predisposed, and relates to disorganization at early stages of psychosis, we investigated glutamate measurements from two ACC voxels in individuals with either genetic high risk (GHR), clinical high risk (CHR) state or with first episode schizophrenia (FES). To avoid the confounding introduced by antipsychotic exposure and cannabis use, we ensured all subjects (including FES, CHR and GHR samples) were drug naive, and none had exposure to cannabis at the time of scanning. We hypothesized that GHR will have substantially lower glutamate levels than both CHR and FES groups; In the symptomatic CHR state glutamate levels will relate to disorganization burden, and in FES where symptoms are more prominent, glutamate levels will normalize due to a secondary disinhibition effect on glutamate tone. Given the variations in the MRS voxel placement in prior studies covering the medial prefrontal cortex, we covered 2 distinct voxels - perigenual and dorsal ACC, that participate in two distinct functional networks that are critical players in the emerging integrative accounts of psychopathology in schizophrenia (Default Mode and Salience Network, respectively)(34) .

## 2. Methods and Materials

### 2.1 Participants

We recruited 96 drug-naïve patients with first episode schizophrenia, 63 clinic high risk, 76 genetic high risk and 67 healthy controls (HC) at Changsha, Hunan, China between 2016 and 2022. Diagnostic assessments for FES patients were completed by two experienced senior psychiatrists, based on the DSM-IV criteria(35). All CHR subjects were screened by Structured Interview for Prodromal Syndromes(36) and fulfilled one of three criteria: Attenuated Positive Symptom Syndrome, Brief Intermittent Psychotic Syndrome or the Genetic Risk and Deterioration Syndrome (GRDS). GHR subjects had at least one first-degree relative with schizophrenia (either sons, daughters or siblings) and did not satisfy GRDS criteria (i.e., were not in a CHR state). All HC had no current or past history of DSM-IV Axis I disorder themselves or a family history of mental disorder. Participants were excluded if they satisfied substance abuse or dependence criteria (including any exposure to cannabis), had any known major physical illness, a history of prior antipsychotic exposure, or head trauma resulting in a sustained loss of consciousness for over 5 minutes or more, any contraindications for MRI.

All participants provided written informed consent to this study, which was approved by the Ethics Committee of the Second Xiangya Hospital (No. S009, 2018) and carried out in accordance with the Declaration of Helsinki.

### 2.2 Clinic assessment

All CHR and GHR participants were assessed positive, negative, disorganization and general symptoms using the Scale of Prodromal Symptoms (SOPS)(36). Symptoms of FES were assessed by Positive and Negative Syndrome Scale (PANSS)(37). Based on a five-factor model of PANSS(38), all FES were also assessed on their positive, negative, and disorganization symptoms. PANSS disorganization factors included summed scores of stereotyped thinking, poor attention, orientation, mannerisms and conceptual disorganization (speech and behavior). SOPS disorganization included summed scores of odd behavior or appearances, bizarre thinking, reduced attention and reduced personal hygiene. All ratings were completed by trained clinicians or researchers who achieved good to excellent inter-rater reliability in PANSS and SOPS use before the start of the study.

### 2.3 Neurocognitive assessment

Given our focus on the ACC, we chose two components of MATRICS (Measurement and Treatment Research to Improve Cognition in Schizophrenia) Consensus Cognitive Battery (MCCB)(39, 40) that invoke ACC engagement (the Chinese version of Stroop Color and Word Test (SCWT)(41) for response inhibition ability and the Continuous Performance Test (CPT) for sustained attention) and two tests that consistently show large effect size reductions in schizophrenia (the Trail Making Test (TMT) for speed of processing and the Hopkins Verbal Learning Test-Revised (HVLT-R) for verbal memory). All cognitive domain scores were converted to T-scores using sample means and standard deviations to the same measurement scale, the overall composite score for global cognition was computed by summing the scores for all four subtests and standardizing this sum to a T-score. Higher cognition scores reflect better cognitive performance.

### 2.4 1H-Magnetic Resonance Spectroscopy

As described in our previous study(42, 43), we measured metabolites using a 3T Magnetic Resonance Imaging scanner (Siemens, Skyra, Germany) with 16 channel head coils at the Magnetic Imaging Centre of Hunan Children’s Hospital. A 10×20×20mm voxel of interest (VOI) was placed in anterior cingulate cortex (pACC) at a ventral and perigenual site (i.e., likely Default Mode Network - DMN site) and a 20×20×20mm VOI was placed in at a dorsal ACC (dACC) site (Salience Network - SN site) (Figure 1). The perigenual ACC site was identified on the anatomical MRI as the cortex directly rostral to the genus of the corpus callosum. We placed the MRS voxel here such that the posterior wall was directly adjacent and did not include the principal medial sulcus. This placement captures a central DMN node, as shown by a detailed anatomical analysis of the functional organization of the cingulate cortex(44). The second voxel was placed on the dorsal ACC (anterior mid-cingulate) region identified as the cortex immediately above the cingulate sulcus, with anterior wall of the MRS voxel lying on an axis that cuts through the posterior border of genu of the corpus callosum perpendicular to AC-PC orientation (as in Dou et al.’s study(45)). A positional query of these voxels on neurosynth.org confirmed Default Mode and Salience Network associations on the meta-analytical maps for the perigenual and dorsal ACC voxels respectively. Every scan was examined post-acquisition and repeated if spectral linewidth was unsatisfactory or changed notably during the acquisition. 1H-MRS spectra used standard point-resolved spectroscopy sequence (PRESS) (svs_se; NEX 80; TR = 3000 ms; TE = 30 ms; spectral bandwidth = 1200 Hz), and pre-saturation pulses of variable power radiofrequency pulses with optimized relaxation delays were used for suppression of the water signal. Water unsuppressed spectra were acquired in the same voxel locations and used the sequence with NEX=8. T1-weighted anatomical images were acquired using 3-dimensional magnetization-prepared fast gradient echo sequences for voxel tissue segmentation (TR = 2530 ms; TE = 2.33 ms; gap = 0.5 mm; flip angle = 7°; FOV = 256× 256 mm; number of excitations (NEX) = 1; slice thickness = 1.0 mm; number of slices = 192).

**Figure 1.**
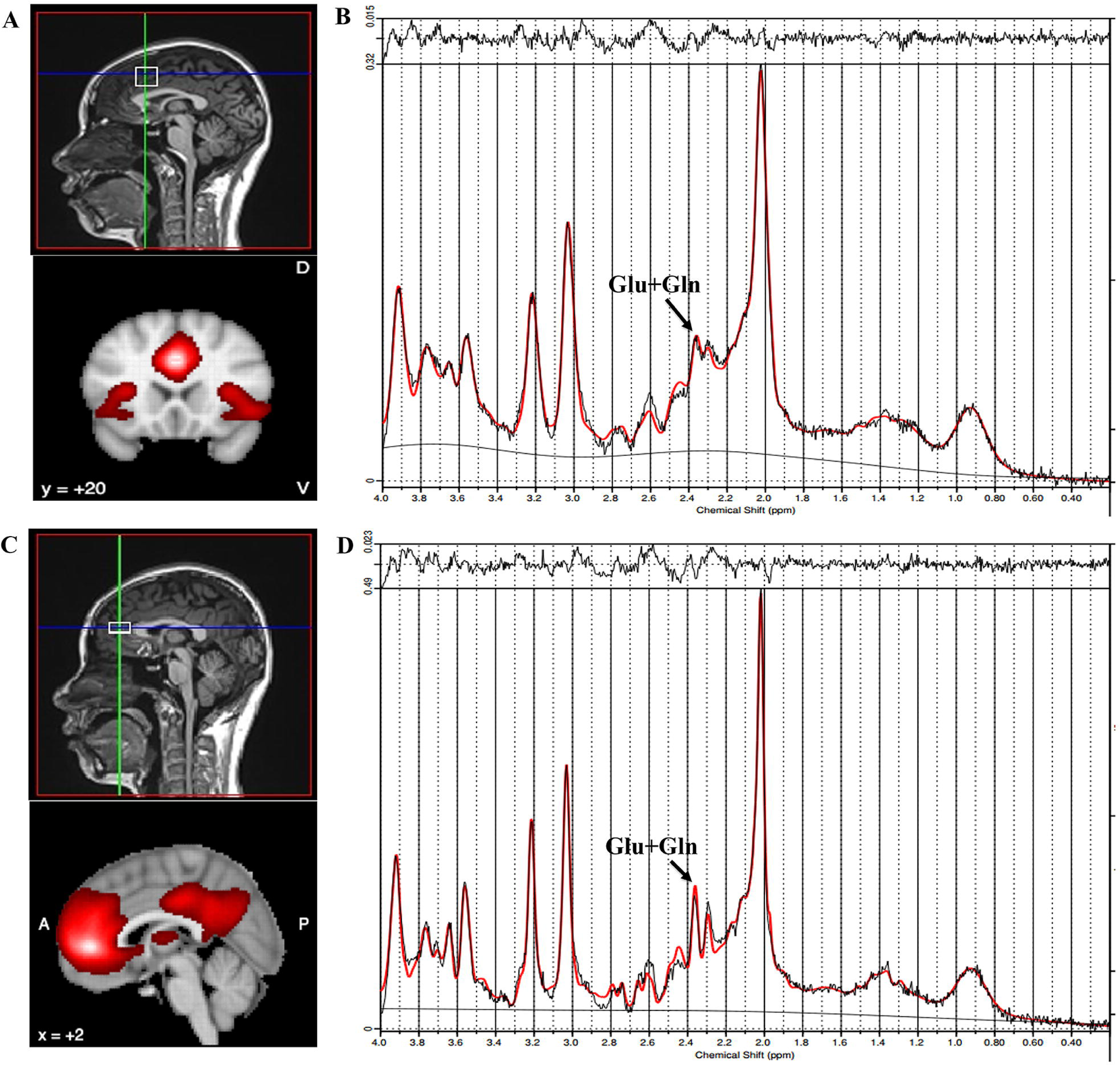
Illustration of voxel placement and example spectrum. (A) Voxel placement in anterior cingulate cortex at a dorsal (Salience Network) voxel. (B) Example spectra obtained from the dorsal anterior cingulate cortex. (C) Voxel placement in anterior cingulate cortex at a ventral, pregenual (Default Mode Network) voxel. (D) Example spectra obtained from the anterior cingulate cortex at a ventral, pregenual voxel. Chemical shift is measured in parts-per-million (ppm).

Analysis of MRI data and quality control are detailed in the supplementary material. The metabolite concentrations were c estimated as the relative concentration of the total creatine (tCr) as the internal reference, due to its relative stability in subjects(46, 47). The primary metabolite of interest was glutamate (Glu); we also analyzed glutamate + glutamine (Glx) in line with the literature to date interpreting Glx as predominantly a glutamatergic signal.

### 2.5 Statistical Analysis

All statistical analysis was completed using SPSS 29 (IBM Corp. IBM SPSS Statistics for Windows, Version 29.0). Demographic variables were compared across groups with the use of General Linear Model and Chi-squared tests for continuous and dichotomous variables, respectively. We used the General Linear Model with pACC and dACC Glu concentration to assess the effect of group and sought between group differences post-hoc using Least Significant Difference correction. A repeated measures ANOVA assessed the effect of group (GHR, CHR, FES, HC) as the between-subject variable, voxel (dACC, pACC) as a within-subject variable. To achieve a normal distribution of symptom scores and to assess the contribution of glutamate to the variance unique to specific symptom domains (rather than overall severity), we calculated the residuals of each syndrome score by adjusting for the overall severity of symptoms from PANSS or SOPS. A stepwise linear regression was used to with glutamate values in either pACC or dACC as dependent variables. We used the following candidate predictors: disorganization, positive, negative symptoms (adjusted for total severity). At each step, variables were added based on R^2^ values with the final model explaining the maximum amount of variance in glutamate. Though the 4 groups did not differ in age and sex, we studied the relationship of Glu and individual symptom severity on these variables using ANOVA (for sex-by-group interaction) followed by t-tests (for sex) and bivariate correlations (Pearson or Spearman as appropriate based on the normality of distributions).

## 3. Results

### 3.1 Demographic, Clinical and Cognitive characteristics

As shown in Table 1, the three groups (FES, CHR, GHR) had no significant differences from HC in age and sex. However, HC has a higher education level than other groups, as expected from the illness-related effects on education outcomes. CHR subjects had a more severe symptom burden than GHR. FES group had the lowest cognitive function among the four groups in CPT, HVLT-R, TMT and composite cognition. HC had better performance than CHR and GHR in TMT, HVLT-R and composite cognitive function, while CHR and GHR were comparable in these tests (HC>CHR=GHR). Both HC and GHR had better sustained attention (indexed by CPT) compared to CHR (HC=GHR>CHR).

**Table 1.** Demographic, cognitions and clinical characteristics of participants. (M±SD) Note: M: mean; SD: standard deviation; HC: healthy control; GHR: genetic high risk; CHR: clinical high risk; FES: first episode schizophrenia; N: number; SCWT: Stroop Color and Word Test; CPT: Continuous Performance Test; TMT: Trail Making Test; HVLT-R: Hopkins Verbal Learning Test-Revised; SOPS: Scale of Prodromal Symptoms; PANSS: Positive and Negative Symptoms Scale.

| Variable | HC(N=67) | GHR(N=76) | CHR(N=63) | FES(N=96) | F/t/$\chi^2$ | P |
| --- | --- | --- | --- | --- | --- | --- |
| Age, years | 20.64±3.49 | 21.55±5.19 | 19.48±4.72 | 21.34±5.60 | 2.52 | 0.058 |
| Sex(male/female) | 39/28 | 38/38 | 36/27 | 52/44 | 1.26 | 0.739 |
| Education, years | 13.94±2.95 | 12.41±3.53 | 11.43±2.71 | 11.79±2.76 | 9.30 | <0.001 |
| Cognition | 54.33±8.40 | 53.06±7.75 | 48.81±9.89 | 44.85±10.05 | 18.30 | <0.001 |
| SCWT | 48.32±8.98 | 49.38±10.25 | 49.38±10.25 | 49.49±9.52 | 1.72 | 0.162 |
| CPT | 54.33±8.40 | 53.06±7.75 | 48.81±9.89 | 44.85±10.05 | 18.30 | <0.001 |
| TMT | 50.47±9.20 | 55.38±4.92 | 49.80±8.44 | 45.73±12.25 | 13.79 | <0.001 |
| HVLT-R | 51.61±6.67 | 56.31±6.21 | 49.35±9.02 | 44.97±11.83 | 21.75 | <0.001 |
| Composite score | 56.82±6.09 | 56.82±6.09 | 49.08±9.14 | 44.43±10.58 | 26.31 | <0.001 |
| SOPS |  |  |  |  |  |  |
| Positive | — | 1.01±1.73 | 10.41±4.91 | — | 14.57 | <0.001 |
| Negative | — | 1.89±3.81 | 11.08±6.25 | — | 10.28 | <0.001 |
| Disorganization | — | 0.72±1.28 | 3.78±2.63 | — | 8.52 | <0.001 |
| General | — | 0.95±1.34 | 4.63±2.91 | — | 9.32 | <0.001 |
| total score | — | 4.56±6.51 | 29.89±10.99 | — | 16.22 | <0.001 |
| PANSS |  |  |  |  |  |  |
| Positive | — | — | — | 21.21±5.88 | — | — |
| Negative | — | — | — | 22.32±7.93 | — | — |
| Disorganization | — | — | — | 21.97±7.51 | — | — |
| total score | — | — | — | 85.02±16.53 | — | — |

### 3.2 Metabolites level

Repeated measures ANOVA revealed that the significant main effect for group (F _(3,298)_ =3.17, p=0.025, partial η2 = 0.031) and voxel (F _(1,298)_ =7.24, p=0.008, partial η2 = 0.024), but there were no group by voxel interaction (F _(3,298)_ =2.23, p=0.085, partial η2 = 0.022). CHR had higher overall glutamate levels compared to other three groups, and pACC voxel had higher levels compared to dACC voxel.

As shown in Figure 2, in pACC, a significant difference was present among the 4 groups for Glu (F _(3,298)_ =2.87, p=0.037, partial η2 = 0.028). The CHR group had higher values than FES (Cohen’s d =0.42) and HC (Cohen’s d =0.35) groups in post hoc tests (Glu: mean (SD) in HC=1.60(0.23); GHR =1.62(0.20); CHR =1.68(0.23); FES =1.57(0.29)).

**Figure 2.**
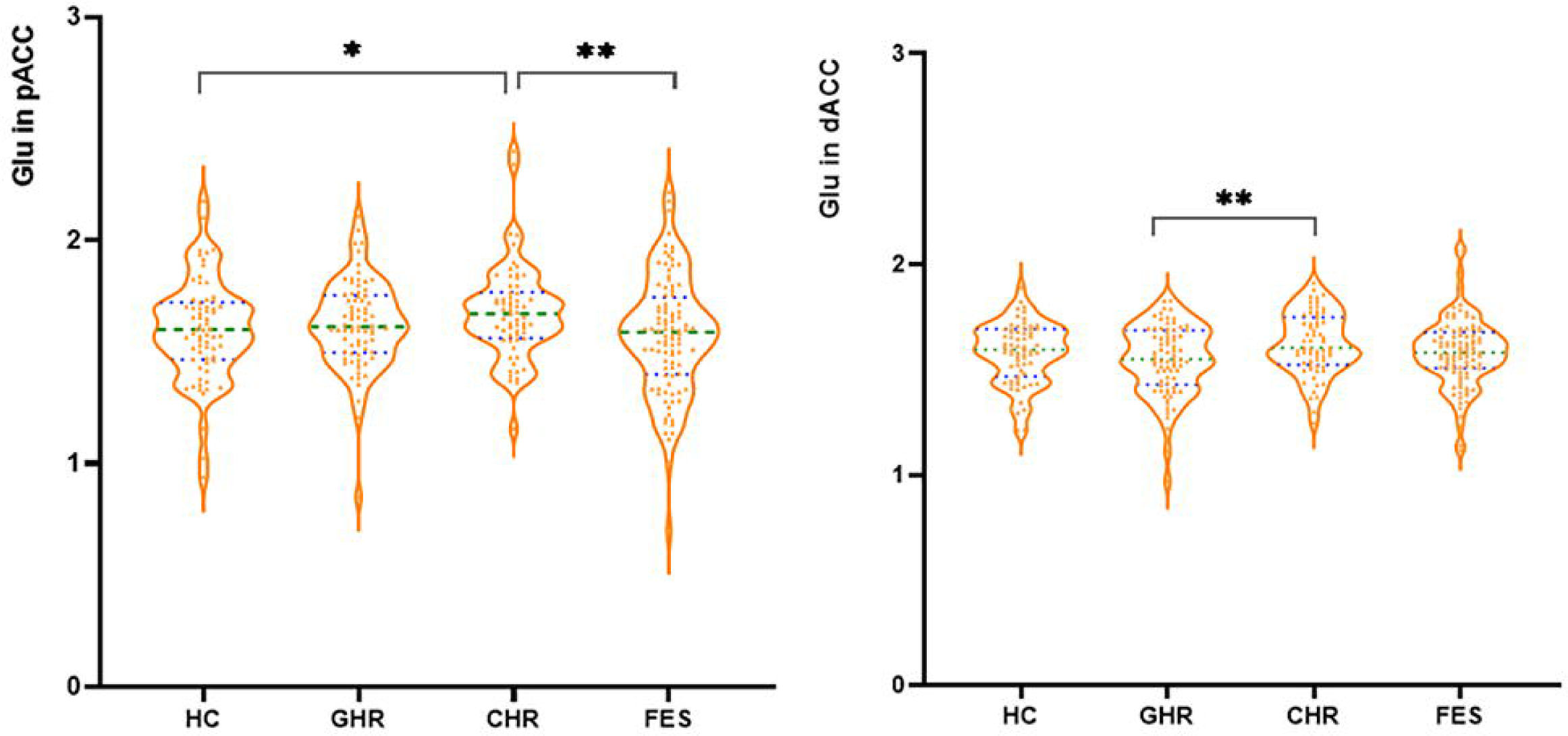
Glutamate metabolite levels in the perigenual and dorsal Anterior Cingulate Cortex voxels. Glu: glutamate; pACC: anterior cingulate cortex at ventral and perigenual voxel; dACC: dorsal anterior cingulate cortex; HC: healthy control; CHR: clinical high risk; GHR: genetic high risk; FES: first episode schizophrenia; The graphs present the individual values (measured in mM), green lines are first and third quartile, and blue line is median. * p < 0.05, ** p <0.01, *** p < 0.001.

In dACC, significant differences were present among the 4 groups in Glu (F _(3,298)_ =2.78, p=0.041, partial η2 = 0.027) levels. GHR had the lowest values for Glu with post hoc tests; especially pronounced when compared to CHR (Cohen’s d= 0.50) (Glu: mean (SD) in HC=1.58(0.145); GHR =1.55(0.17); CHR =1.63(0.15); FES=1.58(0.164)).

The observations from Glx were mostly consistent with the case-control differences reported for Glu, with GHR group showing lowest Glx levels (especially in dACC) (Supplementary Material). Other metabolite values (mean and SD for each of the 4 groups for N-acetyl-aspartate + N-acetyl-aspartyl-glutamate (tNAA); glycerophosphorylcholine + phosphatidylcholine (choline), myoinositol), 1H-MRS quality metrics and voxel tissue volume is also shown in Supplemental materials.

### 3.3 Glutamate levels and symptom

Among the 3 symptom domains, only disorganization predicted pACC Glu in CHR (B=-0.26, Se =0.015, t =-2.08, p =0.042, R^2^[adj] =0.05) and GHR (B=-0.27, Se =0.039, t = −2.43, p =0.018, R^2^[adj] =0.06). In GHR, dACC Glu was also predicted only by disorganization (B=-0.29, Se =0.033, t =-2.58, p =0.012, R^2^[adj] =0.07). Positive and negative symptoms failed to predict Glu levels in both dACC and pACC in GHR and CHR (all p>0.978). The variance of Glu in FES patients was not predicted by any of the 3 symptom domains.

We observed a similar result for Glx models as well, with disorganization in GHR (but not FES or CHR) explaining significant variance in both pACC and dACC Glx; see the supplement for more details.

Correlation analysis (figure 3) revealed that higher disorganization symptoms in the presence of lower Glu (r=-0.26, p=0.042) in pACC for CHR. Higher disorganization was significantly associated with lower pACC Glu (r= −0.26, p=0.026) and dACC Glu (r=-0.29, p=0.011) in GHR group. Other symptoms were not significantly associated with glutamatergic metabolites in pACC and dACC for CHR or GHR. In FES, symptoms did not significantly relate to glutamatergic metabolites in pACC and dACC.

**Figure 3.**
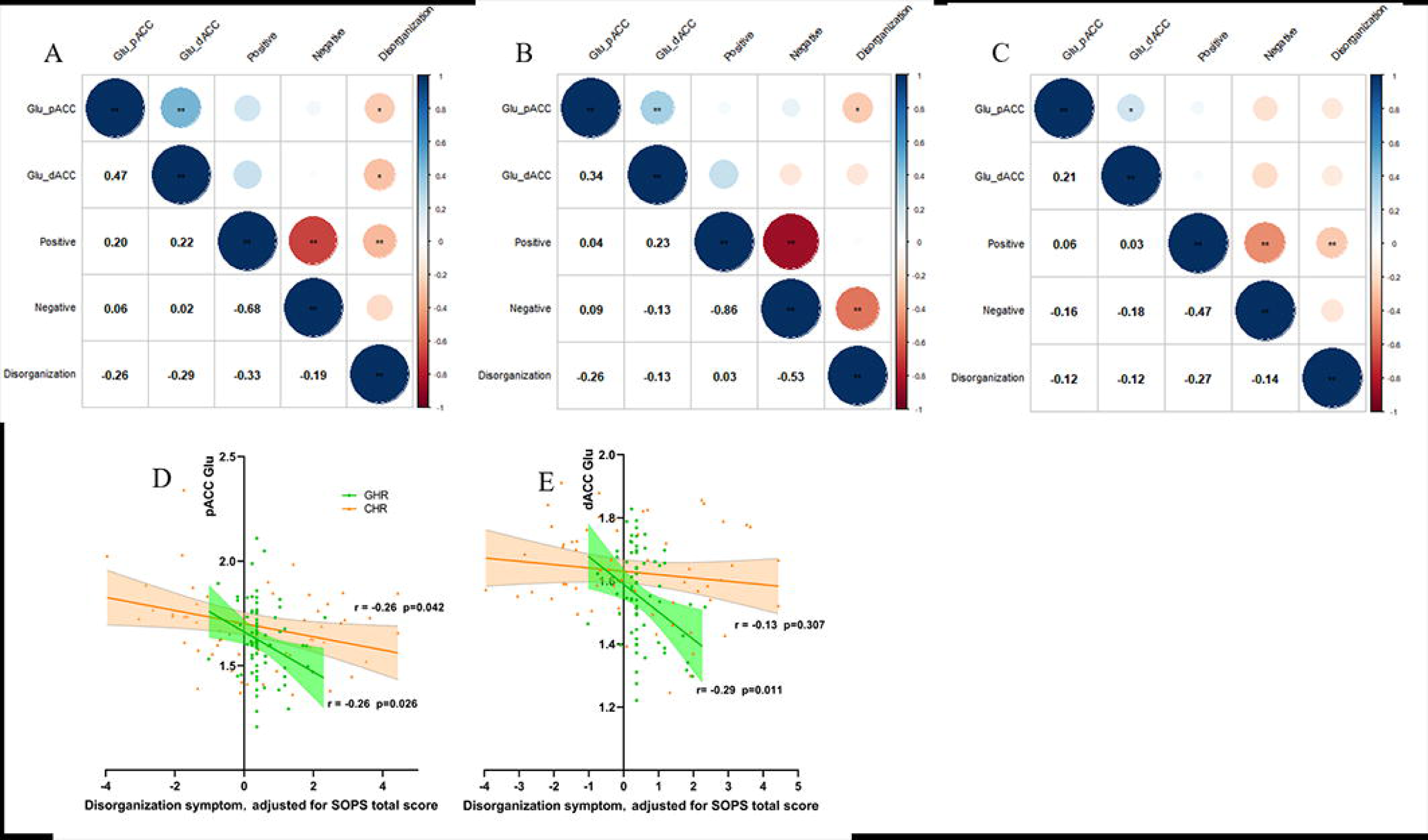
Relationship between glutamate concentrations and symptom after adjusting for total symptom severity for GHR, CHR and FES. A: Heat map of correlation coefficients of SOPS symptom and glutamate in pACC and dACC for GHR; B: Heat map of correlation coefficients of SOPS symptom and glutamate in pACC and dACC for CHR; C: Heat map of correlation coefficients of PANSS symptom and glutamate in pACC and dACC for FES. D: Scatter plot of glutamate in pACC and disorganization symptom for CHR and GHR. E: Scatter plot of glutamate in dACC and disorganization symptom for CHR and GHR. Glu: glutamate; pACC: anterior cingulate cortex at ventral and perigenual voxel; dACC: dorsal anterior cingulate cortex; CHR: clinical high risk; GHR: genetic high risk; FES: first episode schizophrenia.

### 3.4 Glutamate level, age and sex

Age was negatively associated with Glu in pACC (r=-0.21, p<0.001) and dACC (r=-0.30, p<0.001) for the total sample (patients + HCs). Further analysis showed that the negative linear relationship between age and Glu was seen in all 3 patient groups, especially in the dACC (GHR: pACC Glu (r=-0.36, p=0.001); dACC Glu (r= −0.35, p=0.002); FES: pACC Glu (r=-0.21, p=0.042); dACC Glu (r= −0.24, p=0.021); CHR: dACC Glu (r=-0.28, p=0.025)). For HC, age was not significantly associated with Glu in pACC (r=-0.04, p=0.750) and dACC (r= −0.15, p=0.228).

ANOVA analysis revealed that no main effect for sex on pACC Glu (F_(1,294)_ =1.39, p=0.240, partial η2 = 0.005) or dACC Glu (F_(1,294)_ =1.64, p=0.202, partial η2 = 0.006). However, a group by sex interaction was present for dACC Glu (F_(3,294)_ =3.15, p=0.025, partial η2 = 0.031). Further analysis of simple effects revealed that within the GHR group, males had a higher Glu in dACC than females (t=2.93, p =0.004). Higher Glu in dACC was observed in the CHR group vs GHR and HC, in the FES group vs GHR only in females (see Supplemental Figure S4).

## 4. Discussion

We examined MRS glutamate in two voxels of the ACC cross-sectionally in four groups - GHR, CHR, FES and HC. The main findings are (1) Individuals with genetic risk low overall symptom burden had lower glutamatergic metabolites (Glu and Glx) especially in the dACC compared to CHR subjects with emerging symptom burden. Lower glutamatergic metabolites (Glu and Glx) in both dACC and pACC predicted higher disorganization in those at genetic risk. (2) CHR subjects had higher glutamate than other groups in both dACC and pACC voxels, with statistically significant differences emerging in CHR vs. GHR (dACC) or CHR vs. FES and HC (pACC) comparisons. Amidst this group level increase, CHR subjects with higher burden of disorganization still had relatively lower Glu in pACC. Taken together, this supports our hypothetical framework that low glutamate tone reflects the genetic liability (constitutional aberration) that relates to disorganization, while secondary disinhibition that likely elevates/ameliorates glutamate levels occurs at the stage when more prominent positive symptoms start emerging, as in CHR and FES.

Our observation of a null difference in glutamate in both ACC voxels in drug naive FES (n=96) vs HC confirms the observations from the largest drug-naive sample(n=70) to date from a single dACC voxel(48). In FES, several meta- and mega-analyses of medial prefrontal glutamate levels have indicated either no difference from HCs(49) or somewhat lower Glx(50, 51), but this is likely confounded by treatment effects as shown by Merritt et al. mega-analysis who reported a 0.1 unit of glutamate reduction per 100mg increase in chlorpromazine equivalents of antipsychotic dose(51).

Among high-risk individuals (mostly CHR), a previous meta_analysis found that prefrontal Glx was significantly greater when compared to healthy controls(51); we found support for this in our CHR sample in pACC but not in dACC region. McCutcheon et al.(52) expanded Merritt et al. analysis but could not replicate prefrontal Glu elevation after including more GHR subjects (total n=96 genetic high risk subjects included for prefrontal Glu in this synthesis). McCutcheon meta-analysis highlighted the need to separate the symptomatic CHR and the unaffected GHR groups when investigating glutamatergic mechanisms, as some brain regions (especially thalamus) showed GHR specific alterations. In addition, samples included as GHR in MRS studies are generally much older (average age in 40s), by when, the non-occurrence of psychosis reflects resilience, more than the genetic mechanism of the primary pathophysiology. Our GHR sample was matched for age with the other groups, providing a more valid interpretation of the low ACC glutamatergic metabolite levels.

We note that in early stages before the first psychotic break, low glutamate relates to higher disorganization. This is particularly pronounced in GHR sample in whom despite the overall low levels of disorganization, lower glutamate in either pACC or dACC relates to the severity of disorganization specifically, and not to any other symptom domains (positive, negative or total burden). This relationship is restricted to pACC in the CHR sample, where all symptoms are more pronounced, with positive symptoms being particularly more noticeable (∼10 points higher) than the GHR sample.

Contrary to our expectations, we do not find higher dACC or pACC glutamate levels relating to higher positive/negative psychotic symptoms in FES. We expected a glutamate-symptom burden relationship in FES on the basis of a secondary disinhibition in cortical microcircuits resulting from a primary reduction in pyramidal excitatory tone and diminished synaptic gain(53–55). While interindividual variation in symptom severity has not mapped on glutamate differences in ACC in FES in many other studies(56, 57), at a meta-analytic level, studies where samples had higher positive symptoms also reported higher patient-control differences in glutamate levels(8). High glutamate in ACC has been now consistently linked with the status of being a non-responder (58–60) or being treatment-resistant, both indicating persistent high levels of positive symptoms. The lack of Glu-symptom correlation may reflect MRS glutamate measures being an indirect measure for NMDA receptor dysfunction(61), but not the secondary disinhibition that is likely mediated via interneuron downregulation. In fact, dynamic causal models (DCM) of brain connectivity that recovers parameters reflecting interneuron-mediated inhibitory tone, indicates that higher MRS glutamate in dACC relates to higher inhibitory interneuron tone, not disinhibition(62). Interneuron downregulation may be better captured using GABA MRS measures, as shown in a recent MRS-DCM study(63).

Consistent with our previous research(60), we observed that pACC had higher glutamatergic level than dACC, a detailed discussion on this founding can be found in that article. Glu has been now consistently negatively associated with age(8), aligned with our results as well. Schizophrenia is more prevalent in men, with a male to female ratio of 1.4:1(64). Furthermore, males (late adolescence – early twenties) tend to exhibit an earlier age of onset than females (early twenties – early thirties), greater symptom severity, and poorer response to treatment(65, 66). GHR men show secondary disinhibition (higher glutamate) earlier than females, and emerge earlier symptom onset (CHR or FES) resulting from this disinhibitory rebalancing of excitatory and inhibitory transmission, and as a result earlier drop in glutamate occurs in men, which likely leads to similar glutamate level for men in the patient groups. Therefore, the group differences in dACC Glu are primarily attributable to females (see supplement figure S5).

The strengths of our study include the multi-stage sample from GHR to FES stages, scanned at a single site, with two voxels; based on prior meta-analyses, our study sample is one of the largest examining MRS glutamate in GHR as well as drug-naive FES. We also acknowledge several limitations in our study. Firstly, the use of 3T field strength precluded resolving glutamine from glutamate with precision; this resolution has been achieved in several recent studies employing 7T MRS(67, 68). Second, MRS signal reflects the pool of MR-accessible glutamate in the voxel, most of which are not relevant for glutamate neurotransmission. Functional MRS glutamate measures are better suited to provide greater mechanistic information required for this purpose. Third, we lacked longitudinal data on transition/conversion to psychosis in the CHR and GHR samples, limiting the inferences we could make on individual trajectories of glutamate levels in schizophrenia. Fourth, we cannot exclude the possibility of differential glutamate features in other brain regions in GHR/CHR and FES (as shown by Merritt et al.(51), Wood et al.(69) and Fuente-Sandoval et al.(3) in basal ganglia, medial temporal and cerebellar cortex) in schizophrenia.

To conclude, our dual voxel, multistage MRS study without antipsychotic and cannabis confounds supports the presence of a genetically mediated reduction in pyramidal tone in schizophrenia (likely resulting in cortical excitation/inhibition imbalance as per Adams et al. (14)’s hypothesis). This ‘primary hypoglutamatergia’ varies in severity with the degree of subtle disorganization (Meehl’s framework) that is evident prior to the first psychotic break. With emerging psychotic symptoms in CHR and later FES, glutamate levels increase and normalize (likely due to interneuron downregulation that restores cortical excitation/inhibition balance). The putative sequential changes in glutamate levels may explain why the variability of glutamate is higher among patients(70), and why a simple linear relationship between glutamate and symptoms is elusive in schizophrenia. Large, prospective studies on highly resolved glutamate measurements and longitudinal assessment of computed excitation-imbalance are needed for further confirmatory evidence in pursuit of the glutamate-centered mechanism of schizophrenia.

## Acknowledgments

This work was supported by National Key Research and development plan of China (intergovernmental international scientific and technological innovation cooperation project, Grant No:2021YFE0191400), the National Natural Science Foundation of China (NSFC) (Grant No. 81871056 and 82101576), the Fundamental Research Funds for the Central Universities of Central South University (Grant No. 2020zzts287), Science and Technology Innovation Program of Hunan Province (Grant No. 2022RC1040 (ZL)) and China Scholarship Council (Grant No. 202106370192), the Scientific Research Launch Project for new employees of the Second Xiangya Hospital of Central South University. LP acknowledges research support towards this work from the Canada First Research Excellence Fund, awarded to the Healthy Brains, Healthy Lives initiative at McGill University (through New Investigator Supplement to LP); Monique H. Bourgeois Chair in Developmental Disorders and Graham Boeckh Foundation (Douglas Research Centre, McGill University) and a salary award from the Fonds de recherche du Quebec-Sante (FRQS).

## Author contributions

Lena Palaniyappan: Conceptualization, supervision of analysis, writing of original and revised drafts. Ying He, Xiaogang Chen : Supervision, Project administration. Lejia Fan: conceptualization, methodology, acquisition and analysis of data, writing of original and revised drafts. Liangbing Liang: conceptualization. Performed research (acquisition, analysis, and interpretation of data): Zhenmei Zhang, Yujue Wang, Xiaoqian Ma, Liu Yuan, Lijun Ouyang, Ying He, Zongchang Li. All authors contributed to drafting and approved the final version of the manuscript.

