## supplement materials for "A multistage, dual voxel study of glutamate in the anterior cingulate cortex in schizophrenia supports a primary pyramidal dysfunction model of disorganization"

### MRS data analysis

SPM 12 segment tool (FIL Wellcome Department of Imaging Neuroscience, London, UK) was used to segment T1 to extract the gray matter (GM), white matter (WM), and cerebrospinal fluid (CSF) fraction of VOIs. All spectra were analyzed by LCModel version 6.3–1B (http://www.lcmodel.com/lcmodel.shtml), using the standard LCModel basis set, acquired using gamma PRESS at 3 Tesla and 30 ms, and containing below metabolites: L-alanine, aspartate, creatine, phosphocholine, γ-aminobutyric acid, glucose, glutamate, glutamine, glycerophosphocholine, guanidinoacetate, phosphocreatine, L-lactate,myo-inositol, N-acetylaspartate, N-acetylaspartylglutamate, scyllo-inositol, taurine, -CrCH2, lipids (Lip: Lip09, Lip13a, Lip13b, and Lip20), macromolecules (MM: MM09, MM12, MM14, MM17 and MM20), measured in mM unit. The metabolite concentrations were corrected for tissue fraction. Data were excluded if the Cramer-Rao lower bounds (CRLB) exceeded 20% for either of these metabolites. In our dataset, both Glu and Glx spectra fitted with a good degree of precision, with resulting CRLBs averaging around 5% for both, at both dACC and pACC voxels (details in Table S1).

### Glx levels

As figure S1 showed, in pACC, there is no significant difference for Glx (F _(3,298)_ =2.09, p=0.101, partial η2 = 0.021), (mean (SD) in HC=2.06(0.33); GHR =2.06(0.33); CHR =2.16(0.34); FES =2.01(0.40)). In dACC, significant differences were present among the 4 groups in Glx (F _(3,298)_ =3.49, p= 0.016, partial η2 = 0.034), GHR had the lowest values for Glx compared to CHR (Cohen’s d=0.56) and FES (Cohen’s d=0.33), (mean (SD) in HC=1.92(0.23); GHR =1.85(0.26); CHR =1.98(0.20); FES =1.93(0.23)).

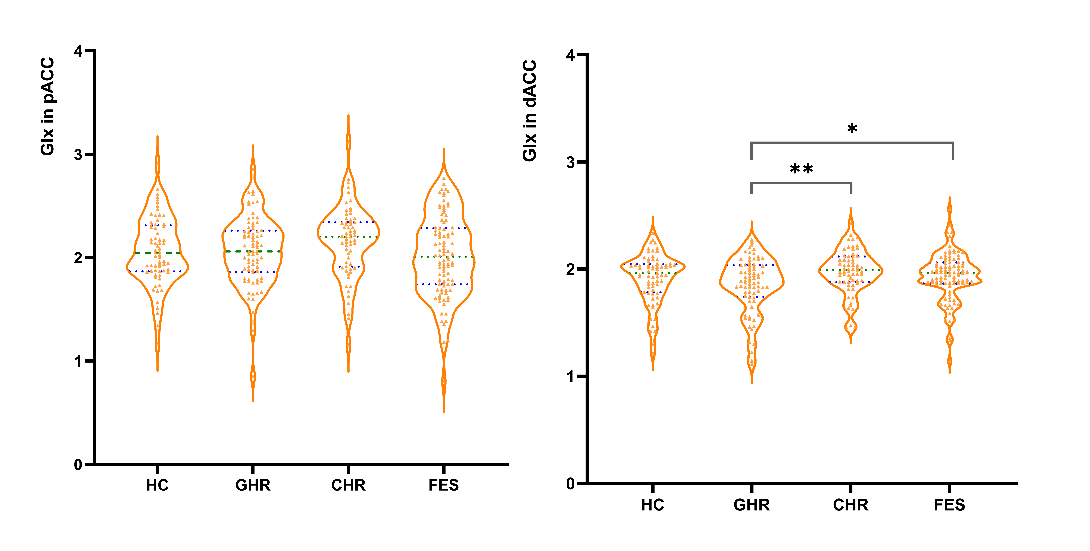

**Figure S1** **Glx metabolite levels in the perigenual and dorsal Anterior Cingulate Cortex voxels** Glx: glutamate + glutamine; pACC: anterior cingulate cortex at ventral and perigenual voxel; dACC: dorsal anterior cingulate cortex; HC: health control; GHR: genetic high risk; CHR: clinic high risk; FES: first episode schizophrenia; The graphs present the individual values, green lines are first and third quartile, and blue line is median. * p < 0.05, ** p <0.01, *** p < 0.001.

### Glx level and symptom

Among the 3 symptom domains, only disorganization predicted pACC (B=-0.28, Se =0.064, t =-2.49, p =0.015, R^2^[adj] =0.07) or dACC Glx (B=-0.35, Se =0.049, t =-3.19, p =0.002, R^2^[adj] =0.11) in GHR. Both positive and negative symptoms were insignificant in predicting Glx levels in both dACC and pACC in both GHR, CHR (all p>0.759). In FES, none of the 3 symptoms were able to predict the variance in Glx levels.

As figure S2 showed, Pearson correlation analysis showed that symptoms did not significantly relate to Glx in pACC and dACC for CHR and FES. Higher disorganization symptom was significantly associated with lower Glx (r=-0.29, p=0.011) in dACC for GHR. Other symptoms were not significantly associated with Glx in pACC and dACC GHR (all p>0.05).

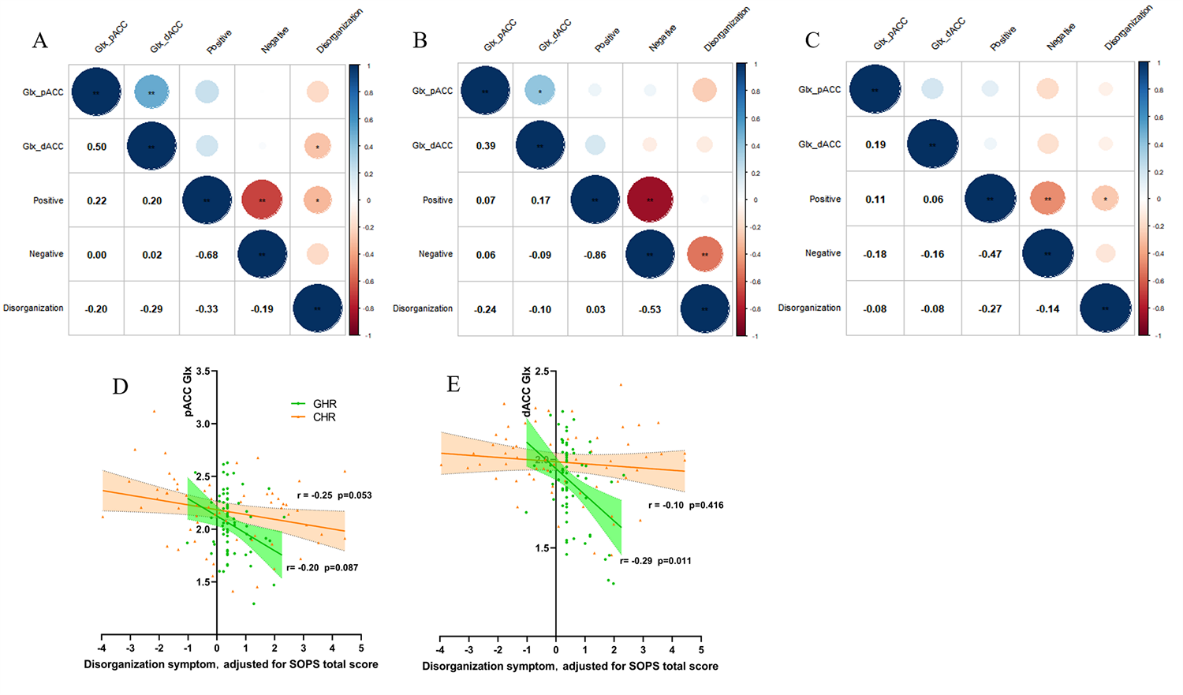

**Figure S2 Relationship between Glx concentrations and symptom after adjusting for total** **symptom severity for GHR, CHR and FES** A: Heat map of correlation coefficients SOPS symptom and Glx in pACC and dACC for GHR; B: Heat map of correlation coefficients SOPS symptom and Glx in pACC and dACC for CHR; C: Heat map of correlation coefficients PANSS symptom and Glx in pACC and dACC for FES. D: Scatter plot of Glx in pACC and disorganization symptom for CHR and GHR. E: Scatter plot of Glx in dACC and disorganization symptom for CHR and GHR. Glx: glutamate + glutamine; pACC: anterior cingulate cortex at ventral and perigenual voxel; dACC: dorsal anterior cingulate cortex; CHR: clinic high risk; GHR: genetic high risk; FES: first episode schizophrenia.

### Other metabolites, 1H-MRS quality and voxel tissue volume

As shown in table S1, we presented information about the concentrations of common metabolites, 1H-MRS quality and tissue volume for four groups.

**Table** **S1. 1H-MRS metabolite concentrations and quality for four groups**

|  | HC(N=67) | GHR(N=76) | CHR(N=63) | FES(N=96) |
| --- | --- | --- | --- | --- |
| **pACC** |  |  |  |  |
| tNAA | 1.10±0.11 | 1.11±0.13 | 1.09±0.13 | 1.08±0.15 |
| choline | 0.22±0.03 | 0.23±0.03 | 0.22±0.03 | 0.21±0.04 |
| myoinositol | 0.91±0.13 | 0.92±0.11 | 0.91±0.14 | 0.93±0.16 |
| Glu CRLB | 5.84±1.50 | 5.48±1.42 | 5.48±1.42 | 6.35±2.28 |
| Glx CRLB | 5.91±1.51 | 5.87±1.50 | 5.62±1.60 | 6.71±2.50 |
| tNAA CRLB | 4.39±1.27 | 5.86±1.50 | 4.89±1.62 | 5.21±1.53 |
| choline CRLB | 4.70±1.34 | 4.50±1.27 | 5.05±1.90 | 5.54±1.82 |
| myoinositol CRLB | 5.01±1.44 | 4.84±1.17 | 5.35±1.99 | 5.65±2.08 |
| SNR | 13.85±4.39 | 13.92±3.40 | 12.21±2.82 | 11.55±3.17 |
| FWHM (ppm) | 0.08±0.03 | 0.07±0.03 | 0.09±0.04 | 0.09±0.03 |
| GMV | 0.53±0.10 | 0.17±0.04 | 0.55±0.10 | 0.52±0.10 |
| WMV | 0.16±0.02 | 0..53±0.10 | 0.16±0.02 | 0.17±0.02 |
| CSF | 0.28±0.10 | 0.29±0.10 | 0.27±0.11 | 0.30±0.11 |
| **dACC** |  |  |  |  |
| tNAA | 1.26±0.07 | 1.21±0.06 | 1.24±0.09 | 1.24±0.11 |
| choline | 0.23±0.03 | 0.23±0.02 | 0.24±0.02 | 0.23±0.02 |
| myoinositol | 0.91±0.08 | 0.89±0.78 | 0.92±0.08 | 0.89±0.08 |
| Glu CRLB | 4.03±0.30 | 4.13±0.71 | 4.08±0.27 | 4.17±1.08 |
| Glx CRLB | 4.58±0.74 | 4.70±1.01 | 4.63±0.68 | 4.83±1.32 |
| tNAA CRLB | 2.54±0.72 | 2.57±0.77 | 2.35±0.60 | 2.45±0.71 |
| choline CRLB | 3.27±0.62 | 3.17±0.70 | 3.05±0.33 | 3.19±0.70 |
| myoinositol CRLB | 3.37±0.49 | 3.38±0.56 | 3.46±0.50 | 3.70±0.91 |
| SNR | 25.97±1.92 | 26.01±2.74 | 26.35±2.31 | 25.08±3.77 |
| FWHM (ppm) | 0.06±0.03 | 0.06±0.03 | 0.05±0.03 | 0.05±0.02 |
| GMV | 0.49±0.07 | 0.23±0.03 | 0.51±0.08 | 0.49±0.07 |
| WMV | 0.24±0.02 | 0.48±0.08 | 0.24±0.03 | 0.24±0.03 |
| CSF | 0.22±0.07 | 0.24±0.08 | 0.22±0.08 | 0.24±0.008 |

Data are presented as mean ± standard deviation. pACC: anterior cingulate cortex at ventral and perigenual voxel; dACC: dorsal anterior cingulate cortex; Glu: glutamate; Glx: glutamate + glutamine; tNAA: N-acetyl-aspartate + N-acetyl-aspartyl-glutamate; choline: glycerophosphorylcholine + phosphatidylcholine. The unit of neurometabolite measurements is mM. CRLB: Cramer-Rao lower bounds; SNR: signal to noise ratio; FWHM: full width half maximum, ppm, parts-per-million; GMV: grey matter volume; WMV: white matte volume; CSF: cerebrospinal fluid.

### Glutamate, Glx concentrations and disorganization items for FES

As figure S3 showed, disorganization item score did not significantly relate to Glu and Glx in pACC and dACC for FES.

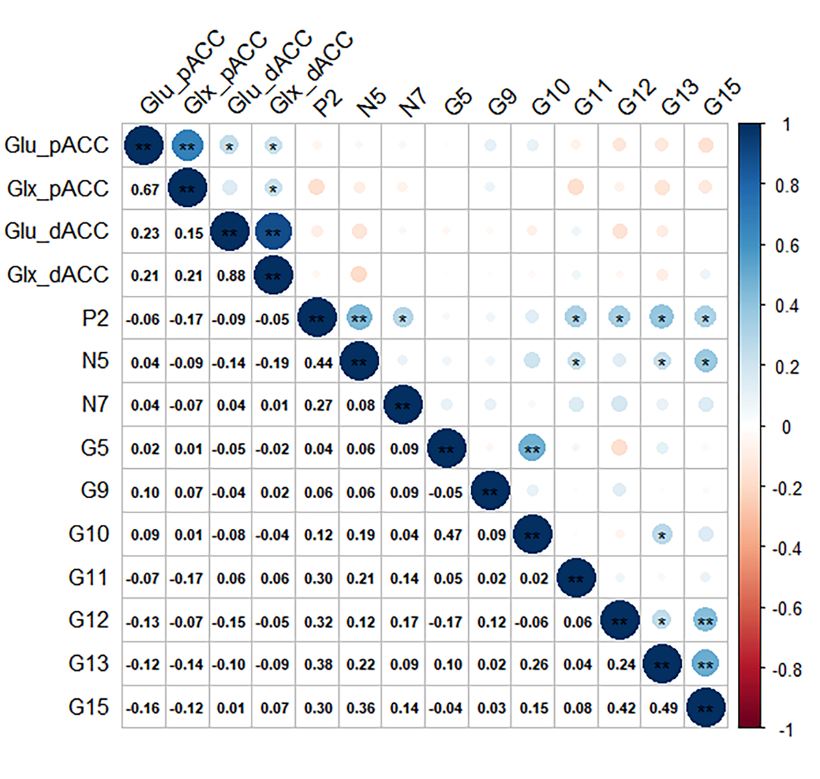

**Figure S3 Relationship between glutamate, Glx concentrations and disorganization items for FES** Glu: glutamate, Glx: glutamate + glutamine; pACC: anterior cingulate cortex at ventral and perigenual voxel; dACC: dorsal anterior cingulate cortex; FES: first episode schizophrenia.

### Glu in dACC for four groups based on sex

.
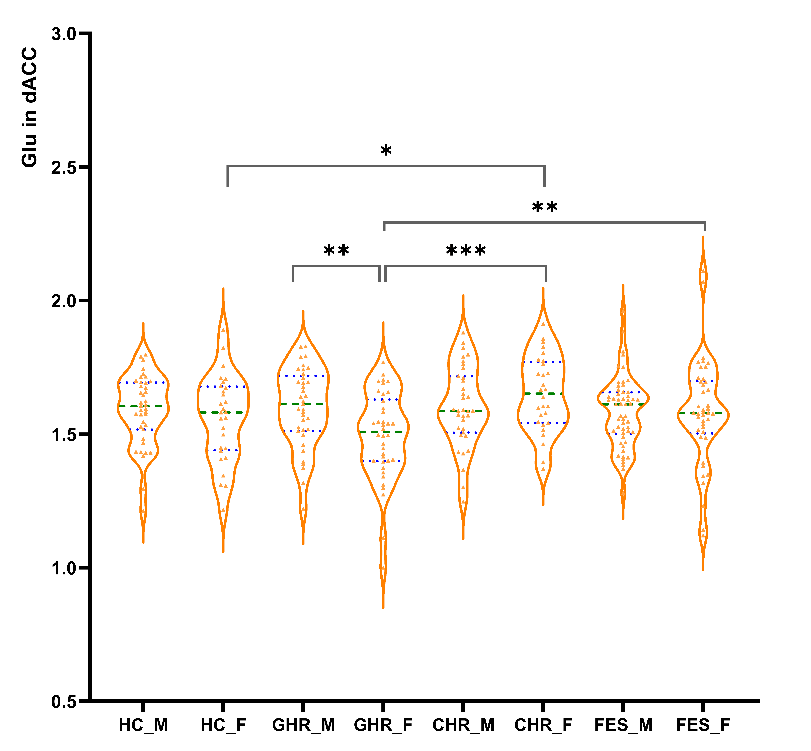

**Figure S4 Glu in dACC for four groups based on sex** Glu: glutamate; dACC: dorsal anterior cingulate cortex; HC_M: male in health control; HC_F: female in health control; GHR_M: male in genetic high risk; GHR_F: female in genetic high risk; CHR_M: male in clinic high risk; CHR_F: female in clinic high risk; FES_M: male in first episode schizophrenia; FES_F: female in first episode schizophrenia; The graphs present the individual values, green lines are first and third quartile, and blue line is median. * p < 0.05, ** p <0.01, *** p < 0.001.

### Hypothesized Glutamate-Symptom trajectories

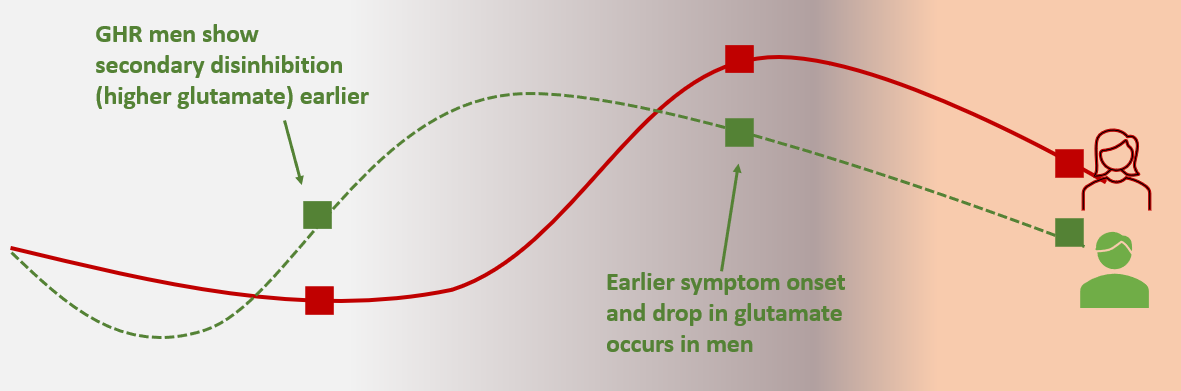
**Figure S5 Hypothesized relationship between glutamate and symptoms of schizophrenia** red line for female, green dashed line for male. GHR: genetic high risk (shaded grey; low symptom burden); CHR: clinic high risk (grey+orange mix, in the middle); FES: first-episode schizophrenia (orange). GHR men show secondary disinhibition (higher glutamate) earlier than females, earlier symptom onset and drop in glutamate occurs in men. Note that the present study does not report on longitudinal data. We make these inferences based on cross-sectional data.
